# Receptor Mediated Delivery of Cas9-Nanobody Induces Cisplatin Synthetic Dose Sensitivity

**DOI:** 10.1101/389122

**Authors:** Philip J. R Roche, Heidi Gytz, Faiz Hussain, Yingke Liang, Nick Stub Laursen, Kasper R. Andersen, Bhushan Nagar, Uri David Akavia

**Author notes:** Corresponding Authors: Dr Philip Roche Dr Uri David Akavia.

## Abstract

The CRISPR/Cas9 system has shown great potential for precisely editing genomic DNA sequences by introducing site-specific DNA cuts that are subsequently repaired by the cell. However, delivery of the CRISPR ribonucleoprotein remains an understudied area and hinders realizing the full potential of the system. We prepared Cas9 ribonucleoprotein complexes chemically conjugated to the 7D12 nanobody and demonstrate receptor-mediated transfection of Cas9 into A549 non-small-cell lung cancer cells via binding to the epithelial growth factor receptor for subsequent cell internalization. We further show that transfection with a Cas9 ribonucleoprotein targeting the BRCA2 gene results in an enhanced sensitivity to the chemotherapeutic drug Cisplatin, and thereby induces a synthetic dose lethality in A549 cells.

## Introduction

CRISPR/Cas9 gene editing has opened therapeutic opportunities that were previously not possible (1, 2). The first wave of CRISPR therapeutic companies (EDITAS, Caribou) and several academic groups focused on *ex vivo* editing (i.e CTX001 for Thalassemia and Sickle cell (3)), and localised injections (CRISPR-Gold in Fragile X syndrome and Duchenne’s dystrophy (4)). Systemic delivery methods of Cas9 include liposomal, cationic polymers, viral and viral-like particles (5); and while Cas9 ribonucleoprotein (RNP) complexes assembled *in vitro* offer higher editing efficiency and lower off-target cleavage than plasmid transfection (6), delivery is generally achieved *ex vivo* by electroporation (7).

Receptor-mediated transfection offers an attractive means to achieve preferential accumulation and increased efficacy of a therapeutic unit (8-11). Recently, small molecule ligands (12) and aptamers (13) have been coupled to the Cas9 protein or a Cas9 containing nanoparticle, respectively, for receptor-mediated uptake into cells. The potential to specifically deliver Cas9 RNP to cells overexpressing a particular receptor type offers many opportunities for targeting the effects of gene editing specifically towards disease-causing cells.

The prospect of using Cas9 for gene knockdown/knockout is clearly understood and widely used, however, applications such as homology directed repair (HDR) using donor DNA templates are still in development in an *in vivo* setting. Having proved to be a powerful screening tool for identifying gene essentiality, Cas9 may be used to overcome chemotherapy resistance (14) and HDR precision would broaden the scope of this application. An ideal Cas9/chemotherapeutic combination would involve targeted delivery through an overexpressed receptor on cancer cells followed by knockout or correction of an oncogene, thereby maintaining or enhancing sensitivity to a small molecule chemotherapeutic.

We chose lung cancer as a model system to evaluate potential therapeutic application. The Epidermal Growth Factor Receptor (EGFR) is overexpressed in lung cancer cells providing an alternative to CD133 for cancer cell identification (15). The Breast Cancer type 2 susceptibility protein (BRCA2) is known to be essential for DNA repair in normal cells (16) and consequently, loss of expression initiates tumorigenesis (17). However, in cancer patients subject to chemotherapy, reversion mutations restoring the open reading frame of the BRCA2 gene can occur and result in resistance to platinum-based chemotherapeutic drugs by maintenance of DNA repair (18, 19). Additionally, it has been shown that knockdown of the BRCA2 gene product by antisense oligonucleotides (ASO) limits cell proliferation in the human lung carcinoma cell line, A549, when co-administered with the cytotoxic drug Cisplatin (20) validating the biological potential for Cas9 knockout studies.

We hypothesize that Cas9 delivered via receptor-mediated transfection can be targeted to BRCA2 creating a synthetic dose lethality of Cisplatin without additional transfection agents. However, the increased protein size with an antibody-targeted nanoparticle systems was shown to impair biodistribution, thereby reducing efficacy, tumour penetration and retention (21, 22). Thus, we chose to use a nanobody (Nb) receptor-mediated transfection system, which maintains target specificity with the added benefit of its smaller size (15 kDa), which does not substantially increase the Cas9 hydrodynamic radius. For our study, we selected two low-EGFR expressing cell lines (A549 and 3T3) rather than A431 (high EGFR expression) as a stringent challenge to the process of receptor mediated transfection (23). Binding of EGFR targeted nanobodies to 3T3 are equivalent to HeLa and A549 cells (24). In this short report, we tested the concept of receptor-mediated transfection of Cas9-Nb complexes leading to a synthetic Cisplatin dose lethality.

## Results

### Nanobodies can be conjugated to Cas9 via NHS/EDC chemistry

To create a Cas9-Nb fusion, we applied amide coupling via NHS/EDC as the simplest method of conjugation. In brief, Nb carboxylic acids were activated by EDC to form the o-acylisourea intermediate that reacts with N-hydroxysulfosuccinimide (Fig. 1A). Sulfo-succinimide forms a stable reactive group in the aqueous phase. Lastly, adjustment of pH to greater than 7.5 improves amide bond formation with primary amines on Cas9, thereby linking the two proteins. The relative simplicity of the chemistry and the few steps required offers flexibility in future applications by allowing for selection of desired Cas9 and Nb variants.

**Figure 1.**
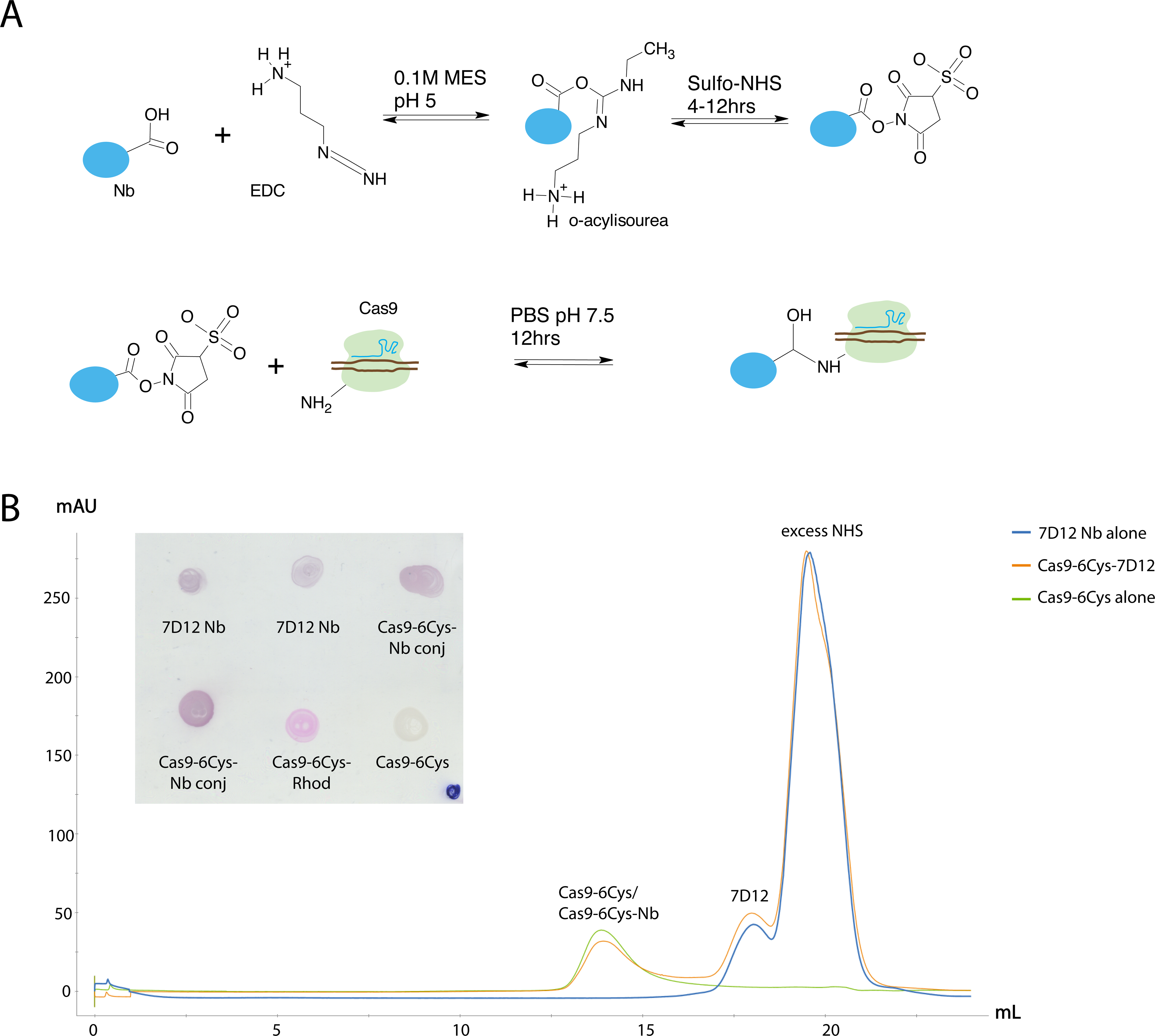
Confirmation of successful Cas9-6Cys-Nb conjugation. **A.** Graphic of the NHS/EDC coupling chemistry **B.** Example of size exclusion chromatograms of Cas9-6Cys, Nb 7D12 and Cas9-6Cys-Nb conjugation. **Inset.** Dot blot was performed to validate that the His-tagged Nb was present in the untagged Cas9-Nb conjugated fractions. Purple color demonstrates the presence of a His-tag and thus the 7D12 Nb. The Cas9-6Cys-Nb conjugated preps give a positive signal, while the control Cas9-6Cys-RHOD (pink) and unlabelled Cas9-6Cys (white) are negative for the His-tag.

To explore the potential for Cas9 nucleases to enter the cell by receptor-mediated transfection, we developed two nuclease variants. The first was Cas9NLS fused to a monoavidin domain (Cas9MAV). This nuclease is desirable for future homology directed repair (HDR) experiments using biotinylated donor DNA. The second was Cas9NLS with 6 C-terminal cysteines (Cas9-6Cys) to facilitate improved protein labeling. Bovine serum albumin (BSA) was included as a control and all proteins were labelled with thiol coupling of the tetramethyl rhodamine (RHOD) fluorophore. The above described Cas9 variants were chemically conjugated to the EGFR nanobody, 7D12 (25) to form Cas9-6Cys-Nb and Cas9MAV-Nb, respectively.

Fig. 1B shows the comparison of three size exclusion chromatograms; Cas9-6Cys, activated 7D12 Nb and the resulting Cas9-6Cys-Nb conjugate. As the elution profiles of Cas9-6Cys-RHOD and 7D12-conjugated Cas9-6Cys-RHOD did not change significantly, a dot blot was performed to validate that the His-tagged Nb was indeed present in the Cas9 fractions upon conjugation. A purple color demonstrates the presence of a His-tag and thus the 7D12 Nb. Control Cas9-6Cys-RHOD (pink) and the unlabelled Cas9-6Cys (white) were negative for the His-tag.

### Non-specific cell penetration of Cas9 and 7D12 mediated Cas9 Transfection

Next, we investigated whether Cas9-6Cys-Nb and Cas9MAV-Nb could target and thus facilitate cellular uptake into A549 NSCLC cells and 3T3 murine cells (Fig. 2A). The experiment compared unconjugated proteins to those with nanobody attachments, and were evaluated at 48 hours post transfection by fluorescence microscopy (RFP channel, Fig. 2B) and fluorescence at 577 nm (Fig. 2A).

**Figure 2.**
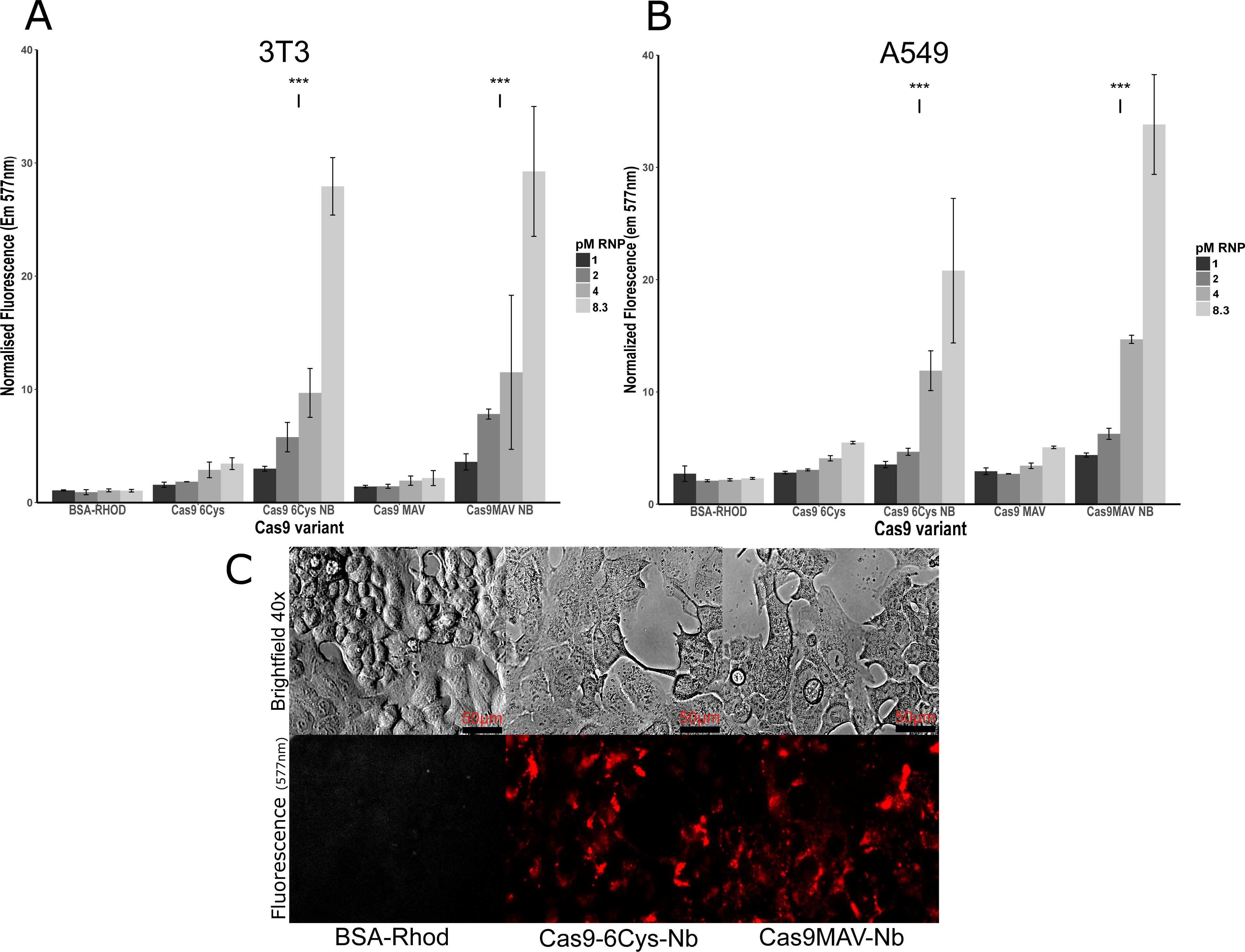
Transfection with 7D12-conjugated Cas9 increases specific cellular uptake of Cas9MAV and Cas9-6Cys. **A.** Dose-response assay of cell penetrating properties of unconjugated and Nb-conjugated protein variants in 3t3 cells. Cells were washed with PBS to remove non-associated labeled protein before cellular uptake was measured in the 96 well plate at 577nm in a SpectraMax M5 plate reader. **B.** Dose response assay in A549 cells. **C**. Examples of transfected cells visualized using fluorescent microscopy.

Cas9-6Cys-Nb and Cas9MAV-Nb showed significant cellular uptake in both cell lines compared to the control BSA-RHOD as well as the unconjugated Cas9 variants. The increase was concentration dependent with **~**30-fold increased uptake at the highest concentration of 7D12-conjugated protein in 3T3, and **~**20-and **~**35-fold increases for Cas9-6Cys-Nb and Cas9MAV-Nb, respectively, in A549. Interestingly, a small non-specific dose response was also observed for unconjugated Cas9-6Cys and Cas9MAV, though overall transfection level was low in comparison to nanobody mediated transfection.

### Cisplatin Synthetic Dose Lethality Assay

To explore the potential for synthetic dose lethality of Cisplatin, we pursued a BRCA2 knockout in A549 cells to enhance the dose response to the drug (Fig. 3). The RNP tested was Cas9-6Cys-Nb complexed to an sgRNA targeting BRCA2. Co-administration of Cas9-6Cys-Nb RNP and Cisplatin was evaluated at a fixed protein concentration (8.3 pmol per well) and varying concentrations of Cisplatin (0.2 - 8 µM Cisplatin). Gene editing is most likely to demonstrate its effect on cell viability post 48 hrs. This was validated at the 24 hr time point where Cisplatin-only and RNP treated cells were indistinguishable (Fig. 3A).

**Figure 3.**
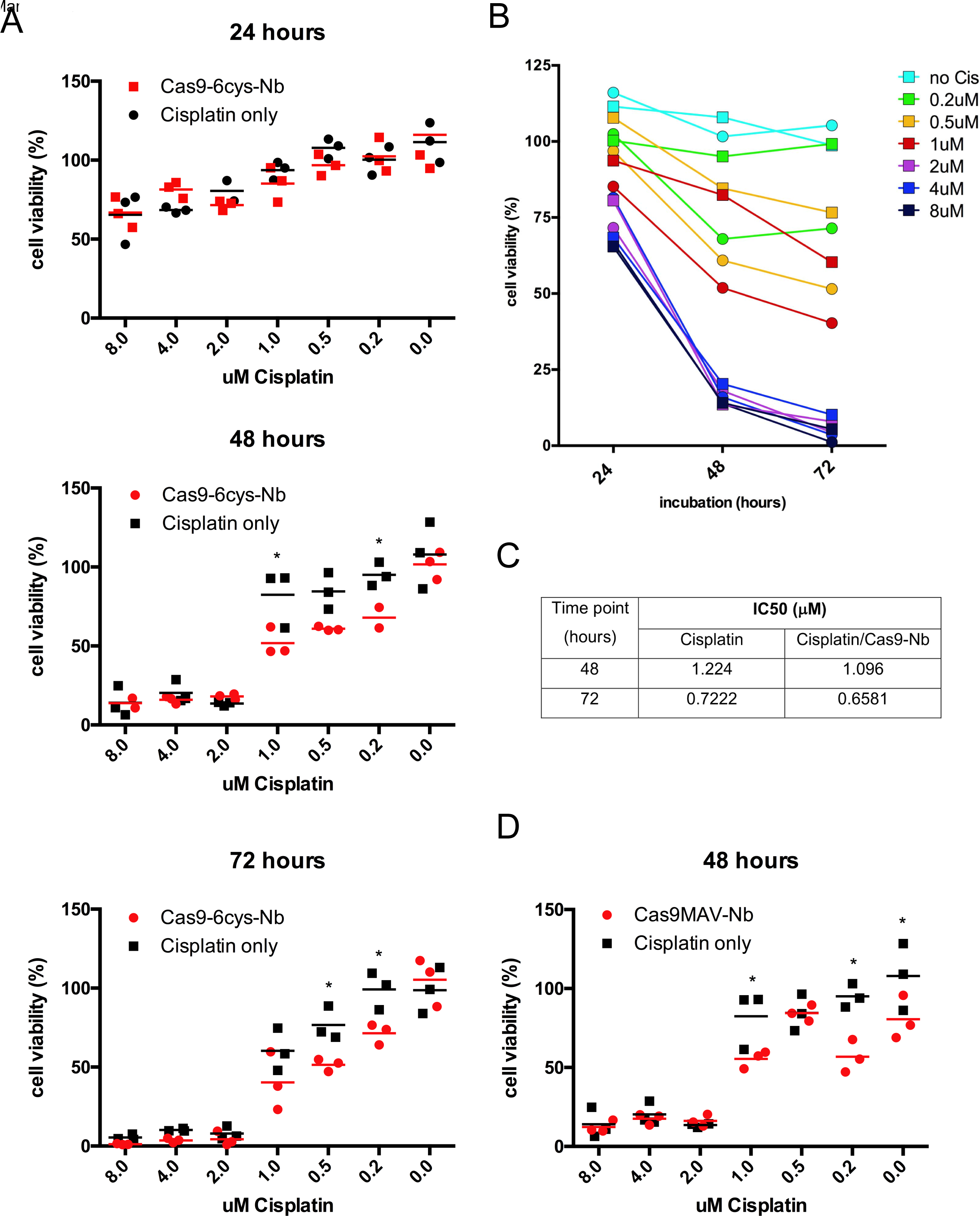
Cas9-Nb conjugates targeted to BRCA2 increase synthetic dose lethality of Cisplatin in A549 cells. **A.** MTT assay was used to measure dose-response of Cisplatin over the course of 72 hours. Percentages cell viability were calculated relative to control Cas9-6Cys-Nb-only (no sgRNA) and untransfected controls. All experiments were conducted in triplicates and significance is denoted by a star, as calculated using GraphPad Prism 6 paired two-way ANOVA. **B.** Summary of mean cell viability percentages at 24, 48 and 72 hours. Cisplatin dose is indicated by colors, while squares denote Cisplatin only and circles denote co-administration with Cas9-6Cys-Nb. **C.** IC50 values calculated on the basis of log(inhibitor) vs. response (three parameters) fitting in GraphPad Prism 6 for 48 and 72 hours, respectively. **D.** MTT assay of Cas9MAV-NB at 48 hours.

Co-administration of RNP and Cisplatin dosage over 72 hrs exposure resulted in the most notable improvement in Cisplatin sensitivity, however, similar significant trends were also observed after only 48 hrs. We know from previous work that 48-72 hrs incubation is sufficient to establish the desired gene edit in a substantial population of cells. Furthermore, Cisplatin-induced apoptosis and cell cycle arrest happens within 8-11 hrs of treatment (26), which is also evident from our results at 24 hrs. Fig. 3B summarizes the development over 72 hrs and shows that the largest fold decrease in cell viability occurs at 0.2-1 uM Cisplatin. Dose response curves were used to calculate approximate IC50 values at 48 and 72 hrs (Fig. 3C), which show that knockout of BRCA2 via Cas9-6Cys-Nb RNP transfection decreases the IC50.

With a view towards future experiments with precision HDR and other targets, we explored whether Cas9MAV behaves similarly and can be delivered to induce synthetic dose lethality. Fig. 3D shows that sensitisation of A549 cells by Cas9MAV-Nb RNP delivery has indeed occurred.

## Conclusion

In this simple proof of concept study, we have demonstrated EGFR receptor-mediated delivery of Cas9 nanobody conjugates and commensurate gene editing leading to synthetic Cisplatin dose lethality. The 7D12 nanobody has a very tight binding affinity (Kd ~0.29 nM) for the EGFR receptor (23). In both cell lines with low EGFR expression, the addition of nanobody resulted in greater uptake of Cas9-Nb conjugates into cells compared to non-conjugated Cas9. Both Cas9-6Cys and our HDR optimized Cas9MAV proteins were successfully delivered and achieved synthetic dose lethality in A549 NSCLC cells. Testing precision HDR with Cas9MAV remains for future work. For therapeutic Cas9 applications, the induction of synthetic dose lethality could be a means to reduce the therapeutic dose and side effects of Cisplatin (27). Furthermore, the well-established NHS/EDC conjugation technique used to fuse nanobody and Cas9 variants of interest, with subsequent purification via gel filtration, brings receptor-mediated Cas9 delivery within the scope of the basic research lab.

There are some limitations of this study: 1) only one sgRNA was evaluated for BRCA2 knockdown where other guides may have generated a higher indel occurrence, 2) the degree of BRCA2 knockout was not characterised by Western Blot leaving the potential for further improvement in lethality, 3) a wider NSCLC cell line screen would be a powerful predictor of the synthetic dose lethality particularly in cells where Cisplatin resistance has developed, 4) recombinant Cas9-Nb fusion proteins were not tested and compared to chemical conjugations and 5) a greater number of biological replicates will eliminate 96-well plate to plate variance. The purpose of this paper is to demonstrate that a biological effect occurs due to Cas9 receptor-mediated transfection. With this limited objective, it is hoped that the principle will be taken by others and more widely applied in enhancing combinatorial Cas9 RNP delivery and small molecule therapeutic studies.

Nanobodies can be generated by selection from recombinant library screening systems (28) or purchased from commercial/academic suppliers with known binding characteristics, rather than consuming significant effort and time in small molecule ligand screening. Additional advantages of this system are the ease of combination with cell/tissue/disease specific sgRNA sequences; the potential to combine multi-valent nanobodies that have enhanced tumour penetration (21) with lower affinity constants (25). Delivery of canonical Cas9 and Cas9MAV make possible gene knockout and high efficiency HDR, respectively, as potential therapeutic modalities to investigate and receptor-mediated Cas9 RNP delivery offers therapeutic opportunities that could be translated into animal models/preclinical evaluations (29, 30). In conclusion, the potential of nanobody-conjugated Cas9 nucleases needs be explored in more depth *in vitro* and *in vivo*, as a means to resolve Cas9 RNP delivery challenge.

## Methods

### Cell Culture and Transfection

3T3 and A549 cell lines were cultured in DMEM supplemented with 10% FBS, 100 U/mL penicillin, and 100 U/mL streptomycin and were maintained at 37°C and 5% CO2. Media, trypsin and FBS were supplied by Wisent. Cells were kept at low passage for experimentation, not exceeding 10 passages before starting fresh cultures from frozen stocks.

For transfections, seeding density was 50000 cells per well (96 well plate) the day before transfection. At a confluency of 60-70%, transfection of RNP was accomplished by addition of 12µl of RNP solution (8.3pmol Cas9 per well), followed by gentle agitation of the plate.

### sgRNA

A single piece sgRNA guide was used in this study. The BRCA2 sgRNA (protospacer sequence GCAGGUUCAGAAUUAUAGGG) was designed using Synthego sgRNA designer and synthesised from Synthego with 5’/3’ 2-O-Me ribose and phosphorothioate backbone modifications.

### Nanobody Expression and Purification

The 7D12 nanobody was expressed in BL21 (DE3) cells, induced with 0.5mM IPTG at OD600 = 0.9 and grown ON at 18°C. Cells were either subjected to complete cell lysis by sonication and cleared by centrifugation, or partial lysis to obtain the protein from the periplasmic space, both with similar low yields of 1mg/2L culture. The 7D12 Nb was subjected to Ni affinity chromatography in buffer A (50 mM Tris pH8, 500 mL NaCl, 5% glycerol, 1 mM PMSF, 20 mM imidazole) and eluted with 500 mM imidazole before being purified by gel filtration in 20 mM HEPES pH 7.5, 150 mM NaCl.

### Purification of Cas9 proteins

SpCas9 fusion constructs were expressed in BL21(DE3) Rosetta2 cells grown in LB media at 18°C for 16 h following induction with 0.2 mM IPTG at OD600 = 0.8. The cell pellet was lysed in 500 mM NaCl, 5 mM imidazole, 20 mM Tris-HCl pH 8, 1 mM PMSF and 2 mM B-me, and disrupted by sonication. The cleared lysate was subjected to Ni affinity chromatography using two prepacked 5 mL HisTrap columns/3 L cell culture. The columns were extensively washed first in 20 mM Tris pH 8.0, 500 mM NaCl, 5 mM imidazole pH 8.0, 2 mM B-me, followed by 20 mM HEPES pH 7.5, 200 mM KCl, 10 % glycerol, 0.5 mM DTT, before elution with 250 mM imidazole. The His-MBP tag was removed by overnight TEV protease cleavage w/o dialysis. The cleaved Cas9 protein was separated from the tag and co-purifying nucleic acids on a 5 mL Heparin HiTrap column eluting with a linear gradient from 200 mM - 1 M KCl over 12 CV.

Gel filtration of Cas9 proteins and Nb conjugates were performed on a Superdex 200 increase column in 5% glycerol, 250 mM KCl, 20 mM HEPES pH 7.5. Eluted proteins were concentrated and stored at −80°C.

### Fluorescent Cas9-Nb Conjugations and Nanobody Biotinylation

Cas9 proteins used in nanobody conjugates were fluorescently labelled using maliamide-tetramethylrhodamine, where 4µl of tetramethylrhodamine maleimide (Anaspec, 10mg/ml, 100x molar excess) was added to a 200µl of protein (8-10mg/ml) in degassed Cas9 buffer and reacted overnight at 4^°^C. The reaction conjugates dye via thiol ester formation between dye and cysteines. Purification was achieved using a Pierce dye removal kit (Thermofisher) following manufacturer’s protocol.

7D12 nanobodies and Cas9 proteins were conjugated by a two-step reaction. 7D12 was diluted in 0.1M MES buffer pH 5.5 to 1mg/ml concentration (final volume 500µl) and COOH R-groups were activated using 1-ethyl-3-[3-dimethylaminopropyl] carbodiimide (EDC, 0.5mM final, Geobiosciences) forming O-acylisourea intermediates and the more stable amine reactive intermediate N-hydroxysulfosuccinimide (sulfoNHS, 4mM final, (Geobiosciences).The reaction was allowed to proceed for 4-12 hrs at 22^°^C. The sample was cleaned up by a G-50 micro spin column (Amersham). Amide bond formation occurring between sulfo-NHS and primary amine R-groups of the Cas9 proteins was conducted at 4^°^C overnight (Nanobody and Cas9 variants in 4:1 molar ratio), with pH adjustment to 7.5 using 10x PBS buffer. Complexes were separated from unconjugated Nb and excess NHS/EDC reagents by purification on a Superdex 200 increase column.

### Dot blotting

Briefly, peak fractions from Cas9-6Cys alone, 7D12 Nb alone and Cas9-6Cys-Nb conjugations were dotted onto a nitrocellulose membrane, blocked in 5% low-fat milk, incubated with mouse anti-His tag antibody (1:2000, Biobasic), washed with TBS-T and incubated with anti-mouse IgG, AP Conjugate (1:2000, Promega) before additional washing steps and development with Sigmafast BCIP/NBT (Sigma).

### RNP formation for transfection

20µl of 1x phosphate buffered saline (sterile and 0.22um filtered), 20µl of Cas9 proteins or Cas9Nb conjugates (25 pmol per 3 wells), sgRNA (concentration varied with respect to Cas9 molarity to maintain 1:1 ratio) were combined in a sterile PCR tube, vortexed gently and incubated for 20 minutes at 25°C. 180µl of DMEM was added to each tube and mixed by pipetting, followed by incubation at 37°C for 10 minutes. Serial dilutions were made of the RNP stock for receptor mediated transfection assay.

### Receptor mediated Transfection Assay

96 well plates were seeded with 3T3 and A549 cells. Four RNP concentrations (1 to 8.3 pmol) of each protein (Cas9-6Cys, Cas9MAV and BSA) were prepared from RNP stocks. Each protein was assigned a block of 3 columns (25 pmol total) and each cell line received the 4 concentrations from the RNP serial dilution in triplicates. 7D12-conjugated RNP was introduced and incubated for 48hrs, at which point DMEM media was removed, cells washed with warmed PBS and visualized by fluorescent microscopy (RFP channel) and then fluorescent was read using a molecular dynamics SpectraMax M5 plate reader (Emission 577nm).

### Receptor Mediated Cisplatin Synthetic Dose Lethality and MTT Assay

A549 Cells were plated to 50-75,000 cells per well overnight. For co-administration (RNP + cisplatin), RNP was introduced at 8.3 pmol per well for each cas9-Nb conjugate, followed by immediate Cisplatin serial dilution administration (8 to 0.2µM + DMSO control), with time points of 24, 48 and 72 hrs incubation. Plates were measured at 590nm to be normalised for background. 20 µl of 5 mg/ml MTT was added to each well and incubated for 2 hrs at 37°C. Media was carefully removed and 150 µl MTT solubilisation buffer (40% DMF, 16% SDS, 2% glacial acetic acid, pH 4.7) was added followed by agitation for 1 hour. MTT absorbance was read at 590 nm using SpectraMax M5 plate reader.

### Statistical Tests

The significance of the improvement in dose-response of Cisplatin between Cas9-6Cys-Nb and no RNP was calculated by two-way ANOVA grouped analysis with GraphPad Prism 6 and R.

## Acknowledgements

The authors would like to thank Prof John Silvius for salient advice on the historical development of receptor mediated transfection and the challenges of protein delivery and the Pelletier Lab for equipment and advice.

---

**S1.**
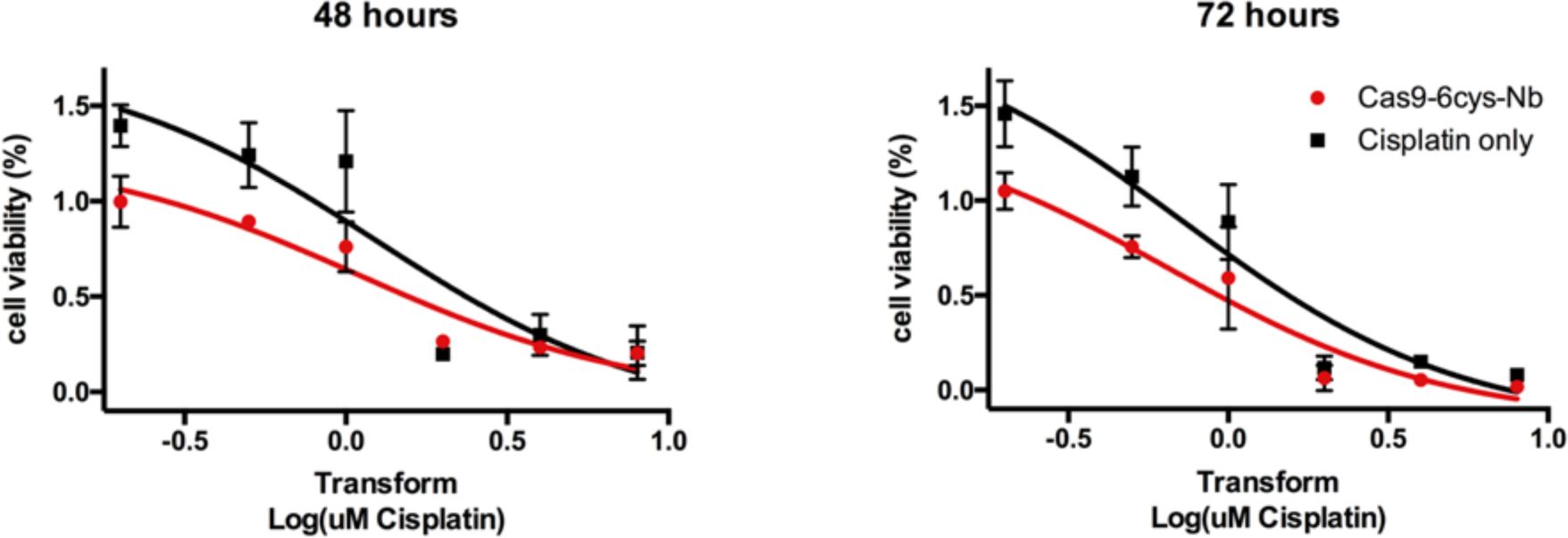
Calculation of approximate IC50 values of Cisplatin. Nonlinear fit of transformed Cisplatin concentrations for determination of IC50 at 48 and 74 hrs, respectively. Analysis was performed with GraphPad Prism 6.

